# Improving peptide-spectrum matching by fragmentation prediction using Hidden Markov Models

**DOI:** 10.1101/358283

**Authors:** Ufuk Kirik, Jan C. Refsgaard, Lars J. Jensen

## Abstract

Tandem mass-spectrometry has become the method of choice for high-throughput, quantitative analysis in proteomics. However, since the link between the peptides and the proteins they originate from is typically broken, identification of the analyzed peptides relies on matching of the fragmentation spectra (MS2) to theoretical spectra of possible candidate peptides, often filtered for precursor ion mass. To this end, peptide-spectrum matching algorithms score the concordance between the experimental and the theoretical spectra of candidate peptides, by evaluating the number (or proportion) of theoretically possible fragment ions observed in the experimental spectra, without any discrimination. However, the assumption that each theoretical fragment is just as likely to be observed is inaccurate. On the contrary, MS2 spectra often have few dominant fragments.

We propose a novel prediction algorithm based on a hidden Markov model, which allow for the training process to be carried out very efficiently. Using millions of MS/MS spectra generated in our lab, we found an overall good reproducibility across different fragmentation spectra, given the precursor peptide and charge state. This result implies that there is indeed a pattern to fragmentation that justifies using machine learning methods. Furthermore, the overall agreement between spectra of the same peptide at the same charge state serves as an upper limit on how well prediction algorithms can be expected to perform.

We have investigated the performance of a third order HMM model, trained on several million MS2 spectra, in various ways. Compared to a mock model, in which the fragment ions and their intensities are shuffled, we see a clear difference in prediction accuracy using our model. This result indicates that our model can pick up meaningful patterns, i.e. we can indeed learn the fragmentation process. Secondly, looking at the variability of the prediction performance by varying the train/test data split, in a K-fold cross validation scheme, we observed an overall robust model that performs well independent of the specific peptides that are present in the training data.

Last but not least, we propose that the real value of this model is as a pre-processing step in the peptide identification process, by discerning fragment ions that are unlikely to be intense for a given candidate peptide, rather than using the actual predicted intensities. As such, probabilistic measures of concordance between experimental and theoretical spectra, would leverage better statistics.

## Abbreviations

DDA: data-dependent acquisition
DIA: data-independent acquisition
FDR: false discovery rate
HMM: hidden Markov model
MS: mass spectrometry
MS1: precursor mass spectrum
MS2: fragment mass spectrum
MS/MS: tandem mass spectrometry
PSM: peptide-spectrum match
TDA: target decoy approach

## Introduction

The use of tandem mass spectrometry (MS/MS) has been pivotal for high-throughput, quantitative analysis of the proteome for discovery experiments. With the iterative advances in instruments over the last two decades, such as improved ion transmission and acquisition rates, thousands of proteins can now be quantified in a single run. Further, techniques such as fractionation prior to the mass spectrometry, longer chromatographic gradients and multiple injections of samples into the spectrometer all increase the total number of proteins quantified. While the exact size and composition of the proteome is subject to spatiotemporal variation, recent studies provide a sense of the bigger picture by analysing the proteomes of many different samples and integrating the results ^1,2^.

As opposed to top-down proteomics, where intact proteins are analysed, in bottom-up shotgun mass-spectrometry approaches enzymatically cleave proteins into peptides prior to the analysis (see ^3,4^ for an overview of MS/MS proteomics). Thus, both the identification and the quantification of proteins in shotgun mass-spectrometry inherently rely on identification of peptides in the sample using their fragmentation spectra. The identification can be achieved in a number of different ways, depending on how much *a priori* information is used in the process. In *de novo* sequencing, peptides are identified by computational analysis of the experimental spectra alone, without the use of any reference. Alternatively, the fragmentation spectra can be compared to a reference, either a library of previously acquired and identified spectra (typically acquired from a similar instrument) or to a database of theoretical spectra for predicted peptides from a selected target protein sequence database, often referred to as *database searching*. While spectral library methods may perform better in theory, their real-life utility depends on the existence and quality of the reference library used^5^, which in turn are built from spectra identified by database searching. Furthermore, an obvious shortcoming of these methods is that they will not, by definition, be able to identify peptides that are not featured in the library. Thus, database searching remains to be a crucial aspect of peptide identification in tandem mass spectrometry workflows relying on data-dependent acquisition (DDA). Data-independent acquisition (DIA) relies on libraries of spectra identified with previously on the same instrument using DDA, in a similar manner as spectral library methods do in DDA setting. In this light, prediction of fragmentation spectra will become even more important as DIA methods become more prominent.

While the specifics of the peptide spectrum matching algorithms vary among search engines ^6^, the fundamental idea is more or less common to all commonly used tools. The standard approach is to filter the target database with the mass of the precursor ion obtained in MS1, then to evaluate for each candidate peptide the similarity between the reference spectrum and the observed spectrum. This similarity is typically evaluated with a scoring scheme that is typically a function of the number of expected peaks from the reference spectra that are observed in the experimental spectra. The best scoring candidates are denoted as a peptide–spectrum match (PSM) with score of the candidate. Given tens or hundreds of thousands of spectra in a dataset, an overall quality measure for the PSMs is ascertained by statistical means, often by controlling the false-discovery rate (FDR), which is estimated using a target–decoy approach (TDA) ^7^. Furthermore, it has been pointed out in literature that compliance to TDA in multi-stage database searches is not entirely a trivial matter ^8^.

Contrary to the thorough statistical methods for estimating overall quality of peptide-spectrum matching process, measures of similarity between reference and experimental spectra tend to be fairly simple as modelling of peptide fragmentation for a complex sample has proven itself to be a difficult endeavour. Despite the differences in the details of various scoring schemes, majority of the widely used database search engines work with the assumption that all fragment ions are equally likely to be observed, when matching a candidate peptide to experimental spectra. However, the assumption that each theoretical fragment is just as likely to be observed is neither well supported nor accurate. On the contrary, for many peptides the fragmentation spectrum is typically dominated by a few daughter ions in the series. Methods that calculate probabilistic scores based on this assumption thus tend to underestimate some candidate peptides and overestimate others, which leads to erroneous ranking of candidates. By improving this step, we can therefore expect to get more identifications at a given FDR cut-off.

For purposes of confident identification, it makes sense to look for fragments that are likely to give a strong signal, and not look for fragments that may be missing or observed at very low intensity. One obvious way to improve the quality of PSMs in database searching methods is to use predictive models of peptide fragmentation, under defined conditions in the instrument. Ideally, a perfect prediction model would bridge the gap between database searching and spectral libraries, by providing a computationally efficient way to compare experimental spectra to a number of candidate peptides. Short of predicting entire fragmentation spectra, a more feasible approach is to predict relative intensities of the daughter ions of a peptide, in a specific charge state. While the additional information provided in this way does not yield any new evidence for a potential match, it does help eliminate negative matches, as well as increase confidence for the positive matches by eliminating fragments that are not likely to be found anyways.

In this paper, we present a method to predict the relative intensities of y- and/or b-ions of a peptide, given its sequence and precursor charge state, based on a hidden Markov model (HMM). With a large body of fragmentation spectra at our disposal, we show that the fragmentation of a given peptide at a particular charge state is highly reproducible, and that methods using machine learning can indeed be used in this context. Furthermore, we demonstrate the robustness of our model with respect to the choice of training data, as well as the performance of our model in comparison to experimental spectra and other fragmentation models. Last but not least, we show the utility of integrating a fragmentation prediction tool into existing peptide identification workflow as a supplement to the existing scoring schemes, rather than replacing them.

## Materials & Methods

### HMM design

Markov models are probabilistic network models, which define a *process* with an inherent state and an outcome associated with that state. Starting from an initial state *S*_0_, the model is in a state *S_i_* at time point *t_i_* and transitions from one state to the another (or possibly the same) iteratively over the time course, in a stochastic manner. In a hidden Markov model, the states in which the model resides at a specific time point are unknown, but the outcomes of the states, called *emissions*, at each time point are observed. Formally defined, a hidden Markov model can be described as a system described by two matrices, *T* and *E*, which represent the transition and emission probabilities, respectively.

We model the fragmentation of a precursor peptide as an HMM, where each fragmentation event (i.e. each b-/y-ion pair) is a process, i.e. a state path, through the model. We define the possible states of the model symmetrically around the fragmentation site, where each state corresponds to one amino acid residue, which are considered to be the emissions (see Figure 1).

**Fig. 1:**
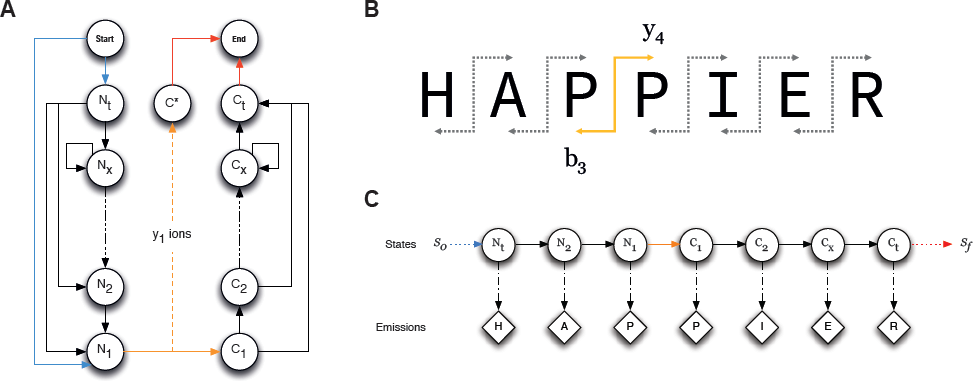
The structure and constraints used for the hidden Markov model. We define an equal number of states (user defined parameter, default 5) to be corresponding to *i* steps away of a cleavage site, towards both ends of the peptide following the peptide backbone (**A**). The states *N_t_*and *C_t_*refer to the ends of the peptides on either side, and *N_x_* and *C_x_* are used as cyclic states to accommodate variable lengths of peptides. A fragmentation event generates an ion pair (**B**), and each ion pair corresponds to a unique state path through the model (**C**).

Additionally, the following constraints are set for the processes: all state paths *i)* start from one of the *N* states, *ii)* must include a transition from *N_1_* to *C_1_* and *iii)* eventually end at state *C_t_* (with the exception of y_1_ ions which end with *C_t_*). The significant number of states on either side of the fragmentation site (*N_1_→ C_1_*) is referred to as the window width, and beyond these are the only recurrent states *N_x_* and *C_x_* which accommodate longer peptides, as well as the terminal states *N_t_* and *C_t_*

This design choice has several significant advantages: first and foremost, each fragment ion pair is guaranteed exactly one state path for a peptide sequence. Thus, during the training of the model for a given precursor sequence and a particular fragment, the state path can be unambiguously determined. Furthermore, this design allows for increasing levels of complexity by expanding the window width around the fragmentation site. Lastly, this model makes no assumptions on underlying biological setting, but rather relies on the electrochemical properties of the peptides. In other words, keeping the choice of protease for cleavage and the fragmentation settings of the instrument constant, the model is agnostic of the sample analysed.

### Model training

Since we aim to predict fragment spectra, the goal is to learn the relative intensities of the fragment ions, given peptide sequence and charge state of the precursor ion. Normally, maximum likelihood estimates for transition and emission probabilities are calculated from a given set of outcomes using the Baum–Welch algorithm. However, given the model structure and the context of the problem, relative intensities can be easily attained from the search results of a shotgun MS/MS experiment, i.e. PSM results from a database search engine can be parsed to extract relative intensities of each identified/matched fragment. Since each fragment yields a unique state path, as per design of the model, once the state path for each fragment is determined, the relative intensity of the ions can be used as a proxy of the probability of following that exact path. In other words, transitions that result in the specific state path of a fragment ion pair can be estimated with the relative intensity of the fragment. Similarly, the relative intensities of fragments are taken as a proxy for the probability of emitting the amino acids that corresponds to the states contained in the state path. In the case of example given in Fig 2C, the relative intensity of y_4_ peak to the other fragments in the y-ion series is used as an estimator of the probability for:

- the transitions: *N_t_* → *N_2_*, *N_2_* → *N_1_*, *N_1_* → *C_1_* …, *C_x_* → *C_t_*, and
- the emissions: *N_t_* ↝ *H, N_2_* ↝ *A, N_1_* ↝ *P, C_1_* ↝ *P*, …, *C_t_* ↝ *R*

**Fig. 2:**
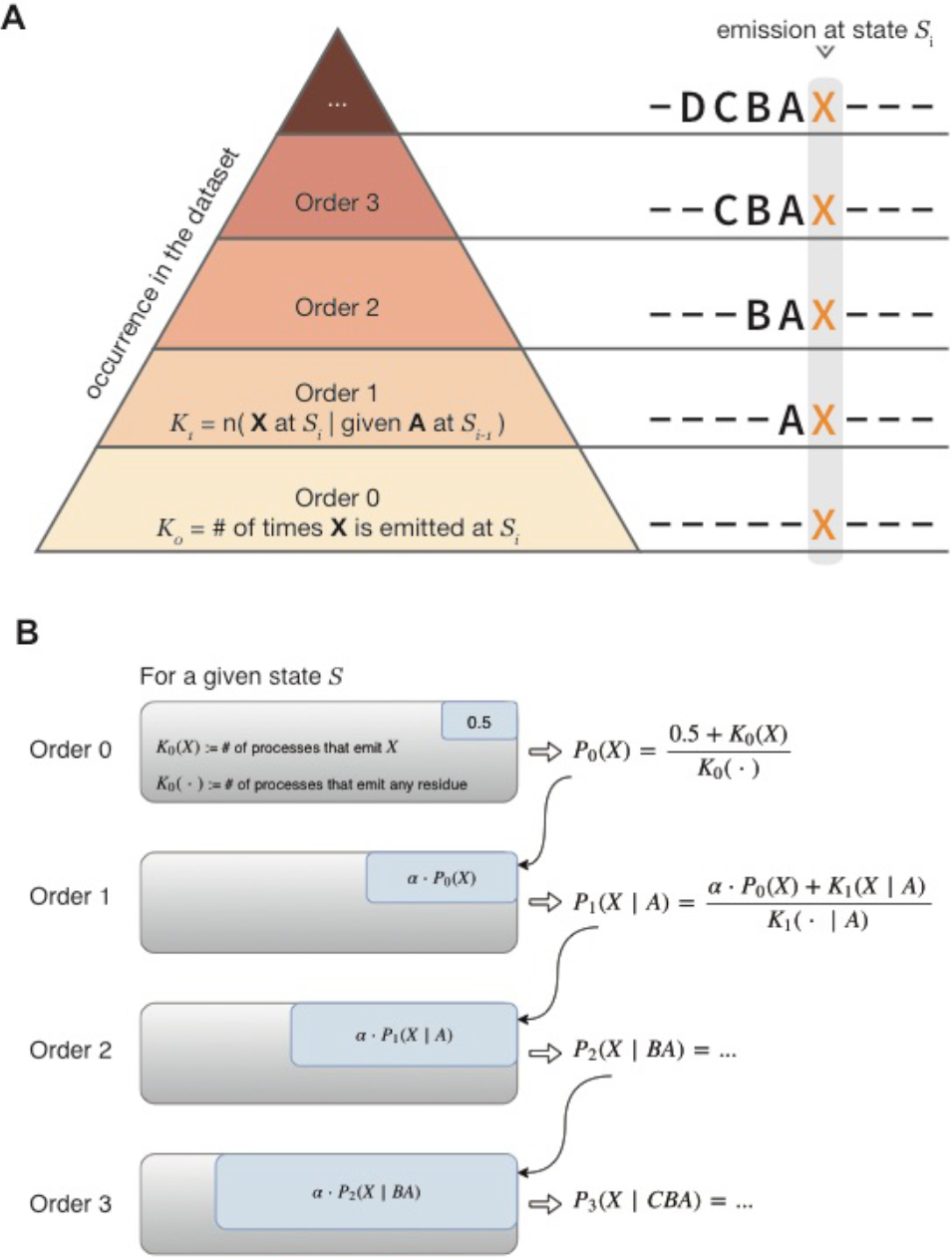
Representation of the model order. A zero-order model only considers emission from a given state, thus it counts the fragments where the unique state path corresponding to that fragment implies a given residue, X, is emitted from that particular state. Higher order models consider the same emission, given previous emissions. It is worth noting that there will be fewer occurrences for each specific case as the model order increases **(A)**. This issue is resolved by adjusting the pseudocounts; a zero-order model is initiated with the pseudocounts = 0.5, in order to avoid absolute zero probabilities, in case a specific emission has never been observed in the training data. For higher order models, the pseudocount is set to the probability of the outcome from one lower order model, weighted by a proportionality factor. **(B)**

As such, the training scheme boils down to processing a large number of annotated spectra (i.e. search results) and accumulating the observed relative intensities of the fragments as well as the fraction of y-/b- ions.

An additional level of complexity is added, however, since the fragmentation of a peptide depends on the length of the peptide and the charge state of the precursor besides the technical aspects such as the type and condition of the instrument and experimental parameters related to the fragmentation process (e.g. collision energy). In order to compensate for any systematic variability, the probabilities are accumulated into different containers for charge states, by default three bins containing +2, +3, +4 or more. Similarly, peptides of different lengths are treated into separate bins, by default 20 bins from 7, to 27+. Both settings are runtime parameters that can be adjusted by the user. Peptides that are singly charged and/or shorter than 7 amino acids are ignored, as they are rarely observed in shotgun MS/MS setting.

### Test data

We have used two different datasets to develop and test the model, in a train-validate-test scheme. We have used the deep proteome analysis of HeLa cells that was published recently ^11^ in order to develop, as well as to test the scalability and performance of the model. This dataset contains over 11 million MS/MS spectra, containing approx. 16 thousand reverse hits, and over half a million peptides from over 14000 proteins.

Furthermore, we a secondary, and much smaller, dataset to show the potential utility of the model. For this purpose we ran our model on HeLa cell lysate, which is used as a standard quality control sample in our lab, analysed in a Orbitrap Q-Exactive HF instrument (ThermoFisher Scientific, San Jose, USA). The raw files were analysed in MaxQuant (ver 1.5.3.8) ^9^ and spectra were searched in against SwissProt human sequence database with 0.01 FDR threshold in Andromeda search engine ^10^. This dataset was used as a primary test of the model. This dataset contains 107283 positive and 98 matching reverse hits, resulting in 31711 peptides identifications.

### Spectra prediction from the model

Once the training data is processed and HMM matrices are estimated, getting predictions from the model is a straight-forward procedure. For a precursor peptide sequence and charge state, first the state path is determined for all y- and b- fragment ions. For each fragment ion, the product of three factors is calculated: *i)* the probability of starting at first state in the path, *ii)* the product of probabilities for the transitions that yield that state path, and *iii)* the product of emission probabilities for the given amino acid sequence. The resulting values are then normalized to 1.0, to yield relative probabilities within the ion series. This process is done separately for y- and b- ions, which allows the user to predict only y- or only b- ion series, as well as both together. In the last case, the y- and b- ion series are weighed using the average fraction of y- ions corresponding to the peptides in the training data with the same charge state and length.

The choice of which ion series to be included in the predictions depends on the experimental setup and ultimately the training data. Since y-ions typically give better coverage when working with HCD spectra ^12^, we have primarily focused on the y-ions in our predictions.

### Model order and estimation of prior weight

Higher order models can be built by considering the emission probabilities to be conditional on the previous amino acids. In other words, considering that the probability of emitting a particular amino acid from a particular state, such as emitting an Isoleucine from state N_3_, might not be independent of the previous amino acid i.e.P(N_3_ ↝ I)≠ P(N_3_ ↝ I| N_4_ ↝ V). We can thus consider *n* amino acids prior to the cleavage site, and build the emission matrix conditionally, such that not only the emission from a particular state are indexed also on the previous *n* emissions for a model of order *n*.

As the model order increases, the probability space for prior emissions increases roughly by a factor of 20, while the probability of observing any specific one of these possible cases in the training data decreases. It is a reasonable assumption that, by default, the probability of emitting Y with prior emissions X_3_X_2_X_1_, is proportional to the probability of emitting Y with prior emissions X_2_X_1_. In other words, when there is a lack of specific outcomes in the training data, the model leverages the probabilities from one lower order model. These default values are typically called pseudocounts and are used to avoid absolute zero probabilities where the training data lacks specific examples.

In order to estimate this weighing parameter, which we refer to as α,we divided training data into slices of equal size and trained a third order model on majority of these slices, then optimized the model using different α values. We used multiple values evenly distributed in log space, (i.e. 1, 2, 4, 8, 16, 32, …, 4096). For each of these models we then checked the correlations of the predicted peak intensities and the experimental spectra in the remaining slices, with data the model has not seen during training (approximately 420000 peptides). We found the 128 to be the optimal value, in terms of average Pearson’s correlation between the predicted fragment intensities and their experimentally measured counterparts. Further analysis of the range between the optimal alpha values in the first run returned negligible improvement for in terms of discovering the optimal value.

### Implementation

We have implemented this model in Python 3.6, the source code including tests and evaluation, is available on GitHub (https://github.com/ukirik/hmsms) together with instructions on how to train a model with custom user data.

## Results & Discussion

### Fragmentation of a peptide in a given charge state is reproducible

Attempting to predict peptide fragmentation is meaningful only if the fragmentation of a peptide is consistent under identical conditions. In other words, without a pattern behind the fragmentation process of a peptide, there is no point in using machine learning algorithms.

In order to address this implicit assumption, we have investigated the peptides with more than 100 fragmentation spectra in our deep proteome dataset, after the quality control steps. We found that the fragmentation pattern of a peptide to be very stable under the same conditions, by looking at the correlation of pairs of randomly selected experimental spectra that were identified as the same peptide at the same charge state (Figure 3). This result indicates that the premise of peptide fragmentation prediction is feasible and meaningful strategy and lays the foundation for our results.

**Fig. 3:**
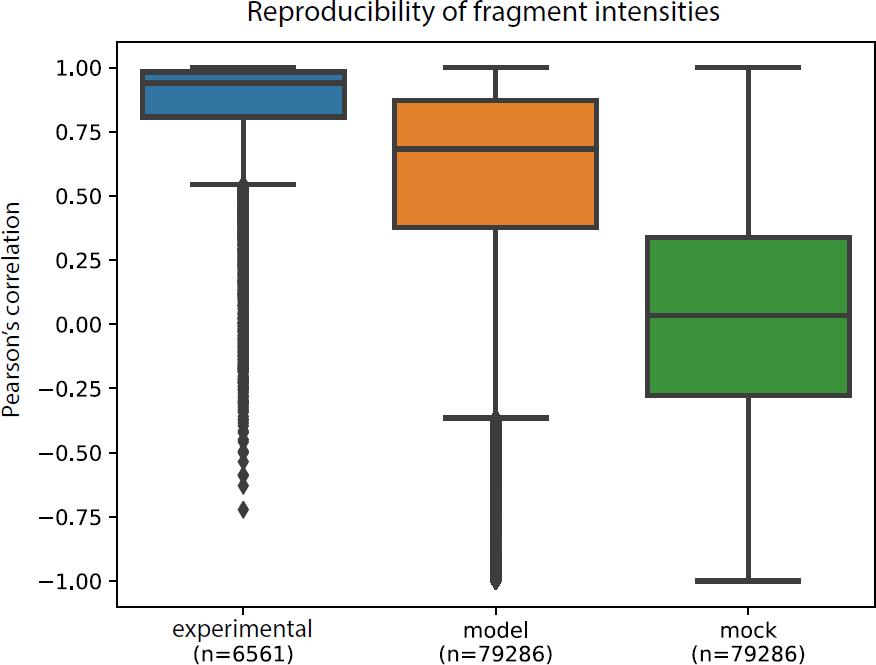
Overall reproducibility, measured as Pearson’s correlation, of the experimental spectra shown together with the prediction performance from the model and a mock model, where the fragments and their intensities are shuffled prior to training. Here we use the reproducibility of the experimental spectra as an upper limit on the performance of a machine learning method, and the mock data as an indicator of null hypothesis. The experimental category consists of randomly selected pairs of spectra for peptides that have been identified with 100 or more spectra.

Furthermore, the overall level of reproducibility we see in our experimental data constitutes a practical upper limit of what can be achieved by prediction algorithms. Using the same metric, we evaluated the accuracy of the predictions of the model is by comparing the predicted fragment intensities against those from measured spectra, where the relative intensity of the fragments from experimental and predicted spectra are the x- and y- coordinates respectively.

### Choosing representative spectra for peptides

Many peptides can be identified with more than one fragmentation spectrum, and the number of spectra available for each peptide will vary considerably within a large dataset. An interesting question in terms of machine learning is thus to determine which spectra to choose for training purposes. This decision is especially important when a collection of datasets is merged into a consolidated training data.

Since we have such a large dataset at our disposal, we set out to investigate how the choice of training data affects the performance of the model. Specifically, with respect to peptides that have been observed more than once at a particular charge state, it is worth considering how choice of representative spectra affects the accuracy of predictions. We considered four different schemes for training the model; for each peptide–charge state pair, we took the best scoring spectrum, the median scoring spectrum, a randomly selected spectrum, and an averaged composite spectrum where intensities for each fragment is averaged out from all available. We trained 4 different models, with all other parameters being the same, based on these different strategies and tested them against 3 different types of test data. We omitted testing against averaged spectra as there is no experimental support for these spectra and may not represent a realistic scenario. Furthermore, in order to give a complete idea of the performance of the model, we provide 4 different metrics: mean Pearson’s correlation with and without missing values, median Pearson’s correlation, as well as a ratio of high correlation to low correlation predictions. The motivation behind the different metrics is explained in the next subsection.

### Evaluation & Performance

How well a model performs in terms of fragment intensity predictions is not a trivial matter. Calculating the correlation between two set of measurements is often the primary approach, however there are several different ways to calculate correlation that consider different aspects of the data points. In this paper, we have considered Pearson’s and Spearman’s correlation coefficients while evaluating the correlation between the normalized intensities of predicted fragments and the observed intensities of the ions. The former evaluates the overall concordance between the pair of values for the same fragment, while the latter evaluates to what extent the internal ranking of the fragments are consistent with each other. We believe that the Pearson’s correlation is the more appropriate measure as the internal ranking of the very low intensity fragments is essentially stochastic, and thus Spearman’s correlation can yield low correlation values for longer peptides with many low intensity fragments.

**Fig. 4:**
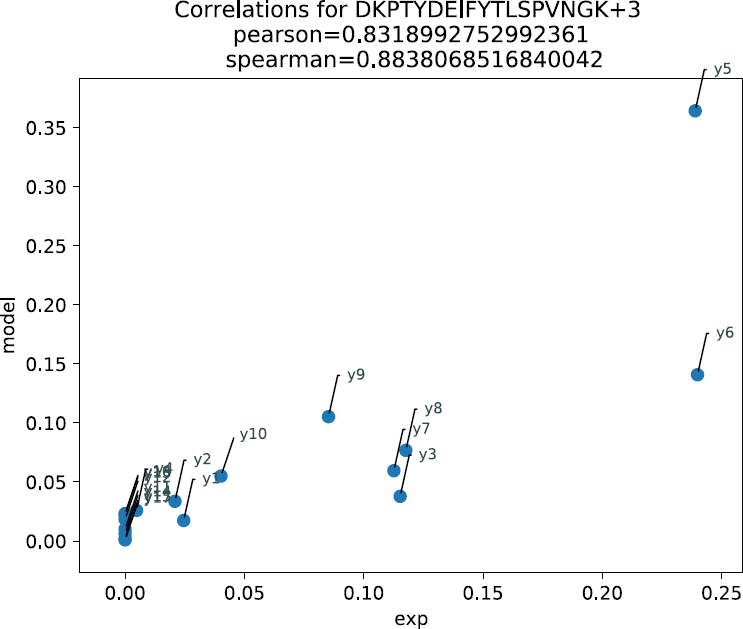
Predictions of the model visualized for an example peptide; fragment y-ions for this peptide are shown in this scatter plot. For each fragment (blue dots) the relative intensity from an experimentally observed spectra and predicted intensity by the model is shown on the x- and y- axis respectively. Pearson’s and Spearman’s correlation is denoted. This example shows the common case where few fragments dominate in terms of relative intensity, several are detected at relatively low intensity and many others are either never identified experimentally or found at negligible intensities.

Additionally, while the use of Pearson’s correlation is appropriate in this context, there are several practical considerations that are of importance. First and foremost, the handling of missing values can vary the performance values substantially. From the viewpoint of a predictor model, each fragment is possible, however some are much more likely to be observed than others. Thus, given a peptide of some length *n*, a probabilistic model will predict some value for all possible y- and b- ion fragments. However, experimental spectra typically have only a subset of the expected ion series identified, for various reasons. Therefore, the experimental spectra will often contain missing values, which cannot be used for the correlation calculation. There are two main ways to deal with this issue; either to exclude the ion fragments with missing values, or to set the missing values to 0, with the implicit assumption that an absence of evidence is evidence of absence. In visual terms, if the fragments were to be arranged on a scatter plot, with their experimental intensities on the x-axis and the predicted intensities on the y-axis, replacing missing values with zero will likely result in a scatter plots, with some points clumping around the origin, and few high intensity fragments far out on the diagonal. If a predictor can get these few intense peaks right, the prediction will be evaluated with a very high correlation value. However, if the missing fragments were removed from the calculation, the resultant correlation coefficient will be much smaller. While neither of the two approaches is technically more correct than the other, it is important to be consistent when comparing results from different datasets, or different models.

Secondly, since the evaluation is done for many peptides, there will be a distribution of correlation values, each coming from one peptide in the test set. The mean of the distribution is likely to be much more susceptible to outliers, i.e. peptides for which the method does exceptionally poor job, than the median. We demonstrate the difference between the different approaches in Table 1. Again, as before, while neither of the methods is more correct than the other, it is important to be consistent in reporting results.

**Table 1:**
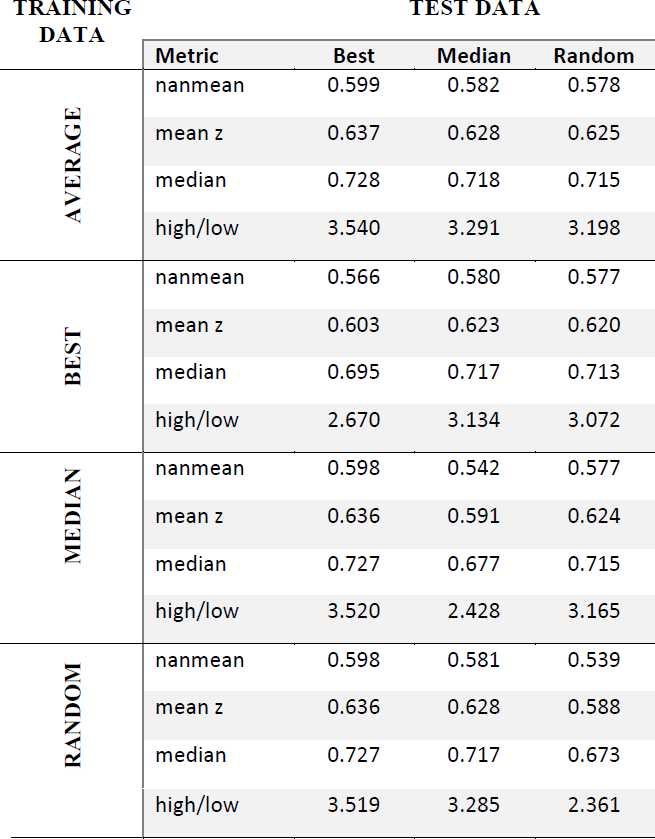
A contingency table showing the variability of performance depending on the choice of spectra used for training vs testing the model. Since the evaluation of predicted ion intensities is not trivial, we provide 4 different metrics; mean Pearson’s correlation with missing values ignored, and set to zero, respectively, median Pearson’s correlation, as well as the proportion of high correlation (> 0.75) to low-correlation (< 0.25) predictions.

| TRAINING DATA | TEST DATA |  |  |  |
| --- | --- | --- | --- | --- |
|  | Metric | Best | Median | Random |
| AVERAGE | nanmean | 0.599 | 0.582 | 0.578 |
|  | mean z | 0.637 | 0.628 | 0.625 |
|  | median | 0.728 | 0.718 | 0.715 |
|  | high/low | 3.540 | 3.291 | 3.198 |
| BEST | nanmean | 0.566 | 0.580 | 0.577 |
|  | mean z | 0.603 | 0.623 | 0.620 |
|  | median | 0.695 | 0.717 | 0.713 |
|  | high/low | 2.670 | 3.134 | 3.072 |
| MEDIAN | nanmean | 0.598 | 0.542 | 0.577 |
|  | mean z | 0.636 | 0.591 | 0.624 |
|  | median | 0.727 | 0.677 | 0.715 |
|  | high/low | 3.520 | 2.428 | 3.165 |
| RANDOM | nanmean | 0.598 | 0.581 | 0.539 |
|  | mean z | 0.636 | 0.628 | 0.588 |
|  | median | 0.727 | 0.717 | 0.673 |
|  | high/low | 3.519 | 3.285 | 2.361 |

In terms of performance evaluation of our model, we first investigated whether or not the model is actually learning anything about peptide fragmentation process. We tested this hypothesis by generating a mock model, i.e. a model that is trained on the exact same training data with one difference, for each spectrum in the training data the fragment ions and intensities are shuffled (y- and b- separately) before updating the emission and transition probabilities. Since the shuffled data should contain no meaningful pattern to learn, we expected the predictions from the mock model to be essentially random, having no correlation with experimental spectra. As expected, predictions from the mock model have practically no correlation and spread all over the spectrum between (−1,1), whereas the corresponding model without shuffling averages around 0.60-0.65 correlation with the experimental spectra. This result indicates that there is indeed learning of the underlying patterns using the HMM model (Fig 3).

After having established that the model does learn from the training data, we set out to estimate how well it performs overall. Since the choice of training data matters, we investigated whether or not predictions vary vastly depending on the train-test partitioning of the input data. We tested this by repeatedly training and validating the model based on different partitions, in a 10-fold cross validation scheme. This comparison not only showed the performance variability based on training data, gave us a good idea about how well the model performs for peptides of different charge states and lengths.

Overall several insights regarding the performance of the model emerged. First of all, we see higher correlations for +2 peptides, compared to +3 and +4 (or more) charged peptides. This finding is not surprising, since having more protons on the peptide backbone increases the chance of having multiply charged daughter ions and thus more complex spectra. Since we collapse the intensities of fragments that are multiply charged, or have neutral losses, onto the single charged ion in the training process, overall abundance of internal ion series or immonium ions are likely to throw off the model.

A secondary insight is that variability is fairly high for shorter peptides with higher precursor charge, +4 or more, and similarly for longer peptides at +2 charge state. One possible explanation for this observation is due to the limited sample size at those specific cases, in other words, that there are comparatively fewer peptides of length between 7-11, and have +4 or more charge, and analogous for longer peptides with +2 charge. Thus, the model typically does not have a lot of training examples for these cases, and similarly the impact one bad prediction (i.e. low correlation between prediction and experimental spectra) is higher on overall variation, if only a few predictions are made, since these cases are equally rare for validation set as in training set.

### Potential utility

As discussed earlier, database search engines come in different flavours and among those, the ones using combinatorial measure of concurrence between theoretical and observed spectra are among the most commonly used ^6^.

Our predictive model can easily be integrated into the existing database search workflows. In the fairly common case where several candidate peptides fall within the mass window of the precursor ion; instead of calculating the probability of matching *k* ions out of all *n* theoretically possible fragments, all of which considered equally likely, our prediction model can be used to discard (or weigh down) unlikely fragments for each candidate. A highly confident peptide-spectrum match should ideally match the high intensity fragments to each other. As demonstrated in Fig. 5, in the case of two candidate peptides with the same number of matching peaks may easily be distinguished, if the intensities of the fragments can be calculated from the sequence alone. Falling short of a perfectly accurate prediction of fragment intensities, we can instead filter, or prioritize, fragments that are more likely to give a strong signal, before matching the peaks.

**Fig. 5:**
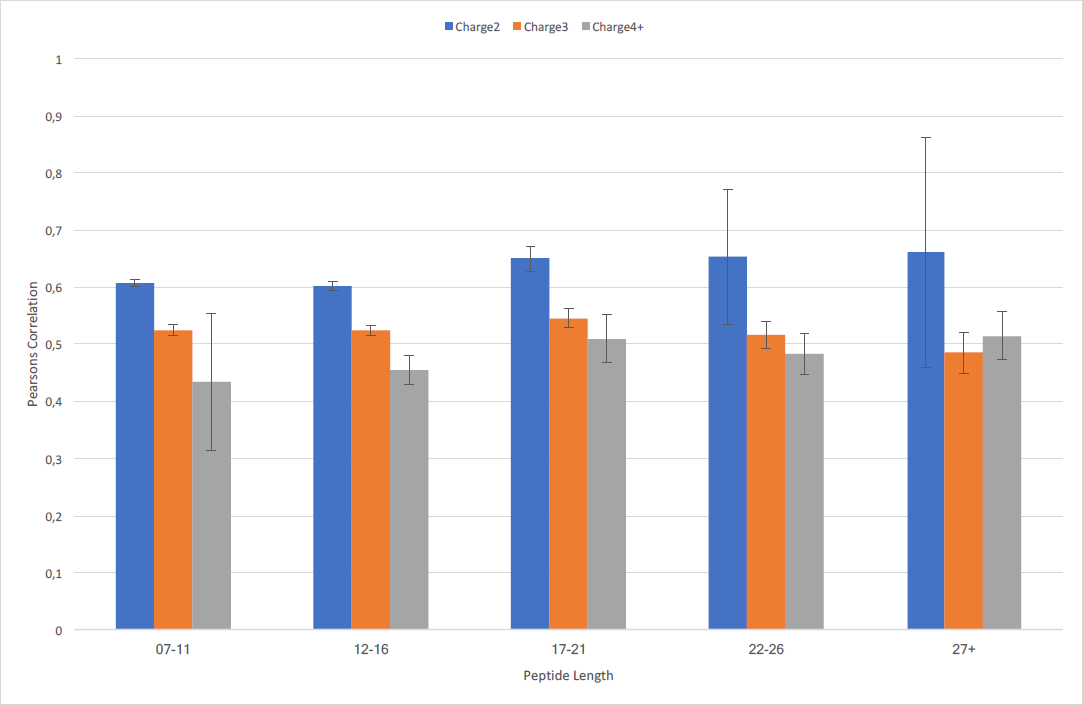
Estimates of prediction variability based on cross validation scheme. For this plot we divided the available data into equally sized partitions and varied the partitions that were used for training and testing of the model.

Figure 6 illustrates how of such a fragmentation predictor can be used in the PSM scoring process. In our example case, we have an experimental MS2 spectrum with two candidate peptides matching the precursor, each with three theoretically possible fragments matching (panel A). Since we do not obtain, or predict, the relative intensities of the candidate peptide fragments to satisfactory precision (panel B), we propose eliminating the lower ranking fragments for the candidates and matching the peaks afterwards. Mathematically this corresponds to decreasing the probability space from matching (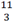) to (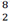) for candidate peptide I, and (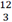) to (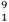) for candidate peptide II. Since these probabilities are in the core of matching candidate peptides to experimental spectra, using this pre-filtering of theoretical fragments should yield improved results, by both decreasing false hits and improving overall confidence in identifications.

**Fig. 6:**
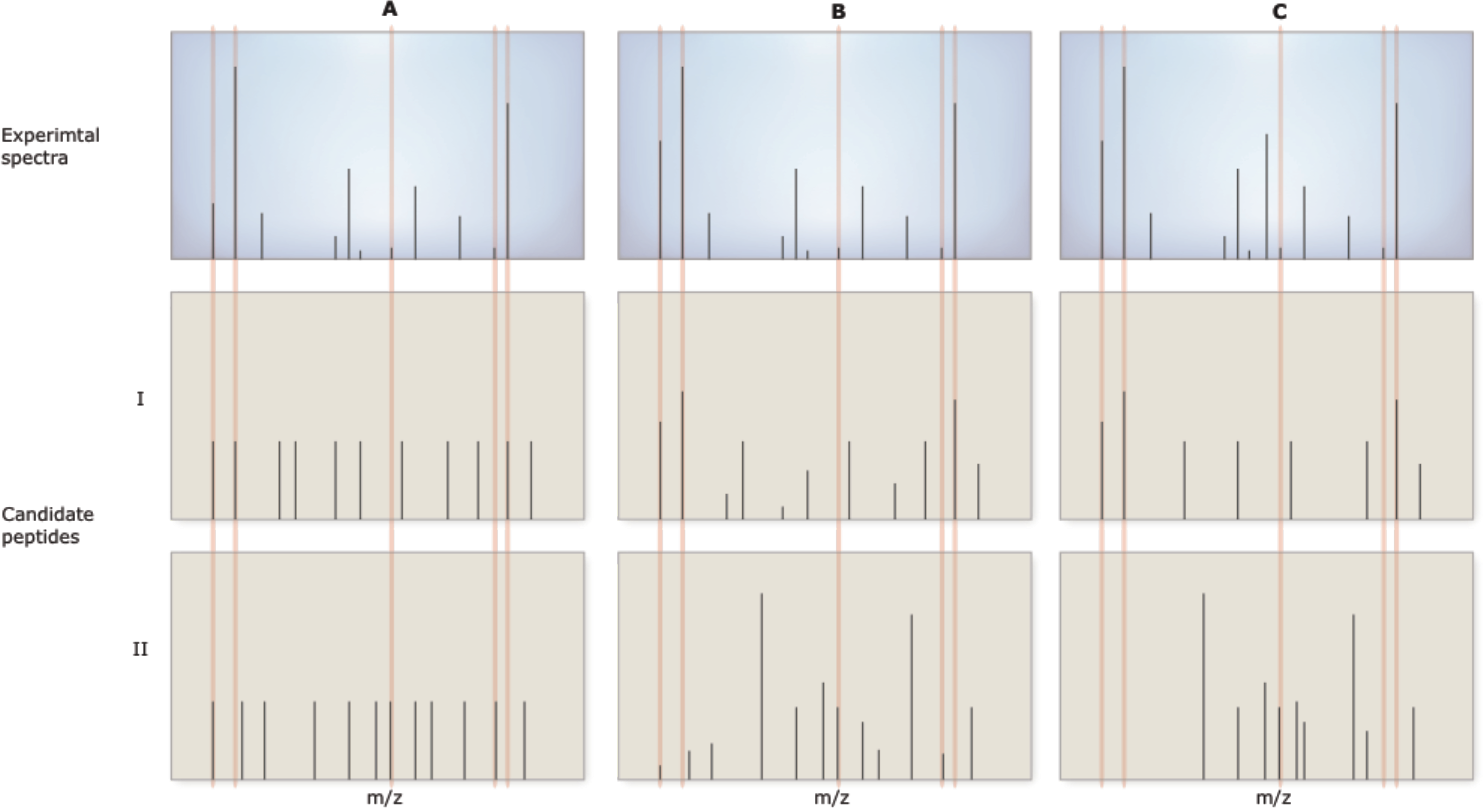
Schematic presentation of the use case of the model. Given an experimental fragmentation spectrum and two candidate peptides that match the precursor mass, the standard database searching procedure (**A**) looks for all fragments giving them the same relative weight. An ideal prediction model (**B**) would be able to predict the relative intensities of the fragments and thus matching ions can be weighted with their relative intensities. Since large-scale accurate prediction of fragmentation spectra has proven to be elusive, we propose a different approach (**C**) where the model is not used to predict the exact relative intensities of the fragments but rather to filter out fragments that are not likely to be strong for a given peptide at a certain charge state.

**Fig. 7:**
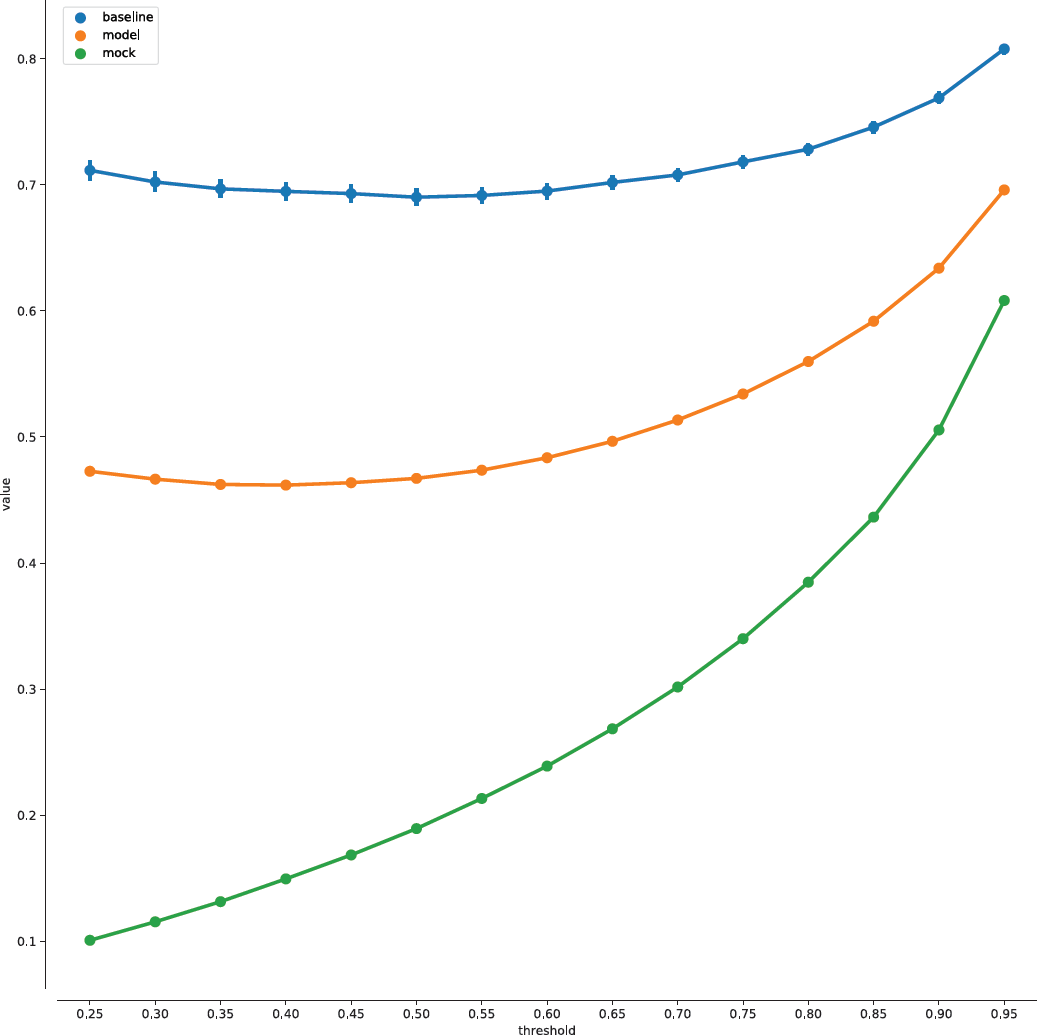
Jaccard’s similarity index can be used as a metric to evaluate the performance of the predictor model, by calculating the proportion of the representative fragment ions. Here the definition of representative fragments is defined as fragments that constitute top *t* percent of the cumulative intensities, given on the x-axis. This figure shows the Jaccard similarity ratios for two randomly picked spectra for each peptide that has been identified more than *N* times (by default = 100) which is taken as an upper limit, similar to the Fig.3. We then look for the similarity ratios for predictions, from an actual model, and a mock model, across different threshold values. In this scheme, *t* in the range (0.6, 0.7) appears to be an optimal value for maximizing the difference between the model and mock, while minimizing the difference between model and the experimental reproducibility level, which constitutes the upper limit for how well a machine learning approach could possibly do.

### Comparison to existing methods

There have been several different approaches previously used for predicting fragmentation in mass spectrometry setting ^13ߝ16^. To our knowledge MS2PIP ^16^ tool, combining XGBoost and random forest algorithms, is the state of the art in terms of prediction of fragment intensities. We have compared our model to MS2PIP, using predictions for 40000 randomly selected peptides from our validation set. For each peptide we picked the experimental spectra with the best score to compare the predictions to the experimental values. Besides the quality criteria used for parsing data out of Andromeda output, these spectra were further filtered to comply with the following criteria: *i)* at least 3 y-ions identified, *ii)* have at least +2 charge, and *iii)* have no more than 1 missed cleavage sites. The peptide sequence and charge state were then submitted as input to the web version of MS2PIP (in batch mode, 10 files) as well as our HMM-model, called HMSMS. Out of the 40000 (peptide, charge state) pairs, MS2PIP returned no y-ions for 5475.

**Fig. 8:**
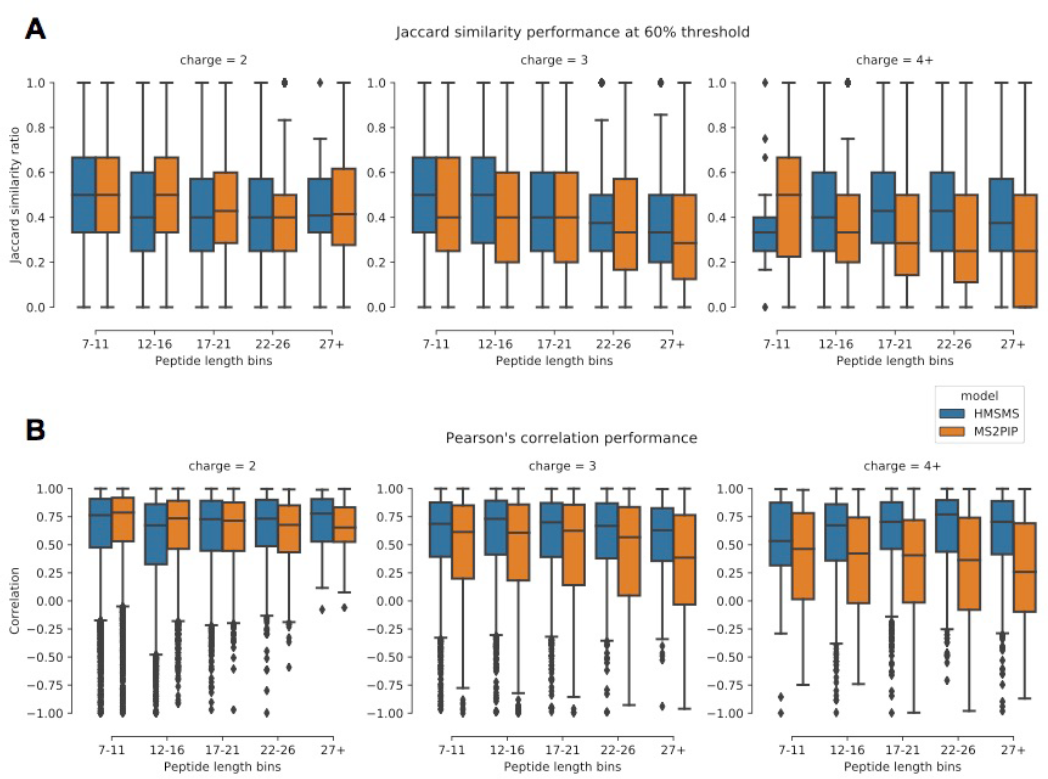
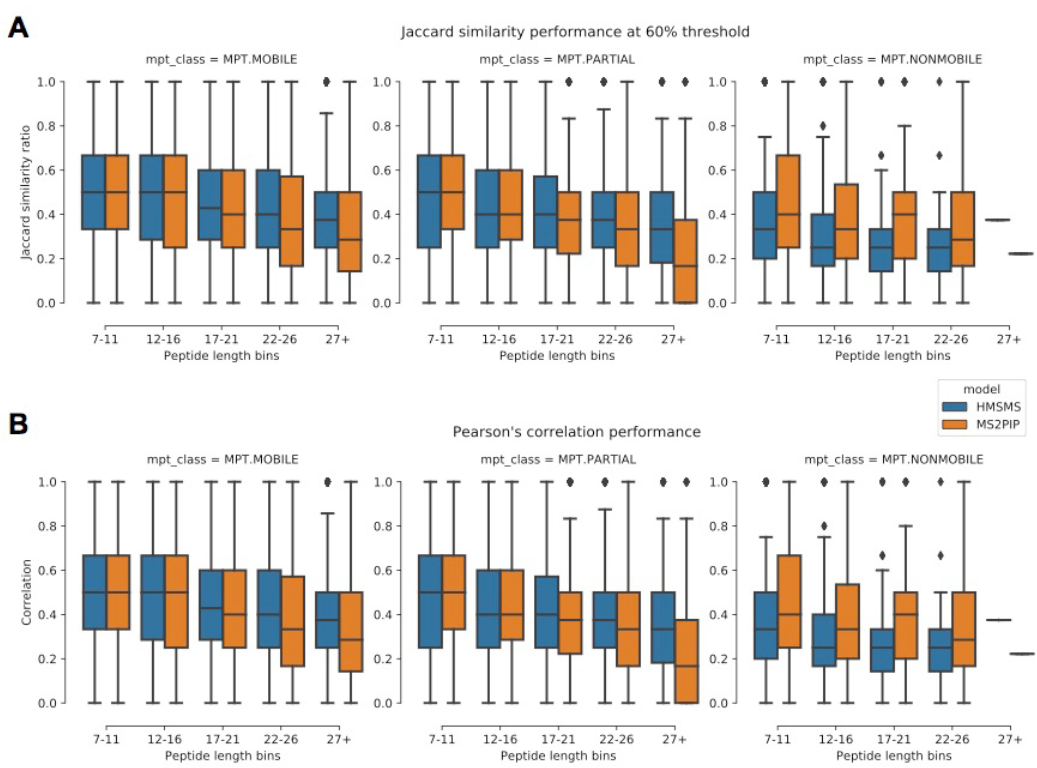
Stratified comparison between MS2PIP and our HMM based model. We have looked at the predictions for 40000 (peptide, charge state) pairs, of various lengths and charge states. The performance of each model was evaluated comparing the predicted fragment intensities versus fragment intensities measured experimentally, using Jaccard’s similarity ratio (**A**) at 60% threshold as well as Pearson’s correlation (**B**).

We have looked at two different metrics, Pearson’s correlation and Jaccard’s similarity ratio at 60% threshold. In both cases, the comparison is between the predictions and the fragment intensities that was measured experimentally. Overall, we found that the methods were comparable in performance, with slight differences in performance in favour of one method or the other. Stratified for different charge states and peptide lengths, we could observe these differences more closely.

We have also investigated whether or not there are other factors that might play a role in the performance of these models. One such factor is the classification of the sequence and charge state pair, in terms of the mobile proton theory. Here, again, we see very comparable results, where MS2PIP appears to do a better job in predicting so called non-mobile peptides, the most difficult class at least in theory, HMSMS has lower variability.

### Computational performance

In terms of computational requirements, our model has very modest demands, both for training the model and predicting spectra. Especially in the case of training, our model relying on the constrained HMM structure is much more light-weight than models that deploy deep learning algorithms.

In the case of our hidden Markov model, even though the training time scales linearly with the amount of training data, the training procedure is of additive nature. Therefore, the training can be parallelized on a multicore machine by slicing the training data into many files and training partial models that are then combined into a final model, in a manner similar to the Map/Reduce algorithms.

As such, training a third order model on approximately 1.6 million spectra using 12 cores takes about 25 minutes. The memory footprint, on the other hand, scales exponentially with model order, due to the expanding space of possible prior emissions. Nevertheless, the model takes approximately 250 MB disk space for persistent storage and approximately 1GB random-access memory during runtime, for a third order model. Once the model is constructed and getting predictions out of the model is on millisecond scale per peptide. Together with correlation analysis, compared to experimental spectra, it is possible to process up to 10-20 peptide-charge state pair per second on a modern (2016 model) laptop running a 2.6GHz Core i7 CPU.

With such modest requirements, the model can easily be deployed on any modern computer and trained on demand unlike models relying on deep learning approaches based on artificial neural networks. We propose research groups to train their own models based on experimental data coming from their own instruments, rather than relying on a model provided by other labs, in a one-size-fit-all fashion. This approach would avoid any disparity between the quality of spectra used for training the model and the spectra on which the predictions will be used. Furthermore, this process can easily be done periodically to keep the models up to date with the state of the instruments. It can even be integrated into the quality control routine for an instrument if a modern computer with a multicore processor is available, which is not practically feasible for models that require specialized hardware and/or take several days to train.

## Conclusions

Peptide identification lies in the heart of the shotgun mass-spectrometry proteomics, and thus it is clear that the reliability as well as the overall utility of the results coming from a shotgun MS experiment will depend on efficient and accurate identification of the analyte peptides. While the instruments have seen many iterative improvements both in hardware and software, the peptide identification algorithms have been more or less constant over the recent years. In this regard, we postulate that peptide identification process can be improved using recent developments in machine-learning algorithms and growing amounts of high quality experimental data available.

Here we have demonstrated a novel approach, based on a hidden Markov model, for the prediction of fragment ion intensities. Our model design features a couple of insightful constraints that take advantage of the nature of the peptides and the fragmentation process, in order to simplify and speed up the training process significantly. More importantly, we show the utility of such a model by using not the actual predicted values but rather as a filtering mechanism for fragments of candidate peptides, such that matching of ions between experimental and candidate spectra is carried out for representative ions. Used in this fashion, our HMM model has the potential to be integrated into existing workflows with little impact on computational time. We provide the source code for the model, as well as all other relevant scripts used for creating all performance comparisons, for the community. In the light of our results, we believe our approach would be a beneficial addition for shotgun MS workflows.

## Acknowledgements

The authors would like to thank David Lyon, Christian Kelstrup, Alex Junge and Helen Cook for valuable input during the development, evaluation of the performance of the model as well as their feedback on the manuscript.

